# Bayesian genome scale modelling identifies thermal determinants of yeast metabolism

**DOI:** 10.1101/2020.04.01.019620

**Authors:** Gang Li, Yating Hu, Hao Wang, Aleksej Zelezniak, Boyang Ji, Jan Zrimec, Jens Nielsen

**Affiliations:** Department of Biology and Biological Engineering, Chalmers University of Technology, SE-412 96 Gothenburg, Sweden; Science for Life Laboratory, Tomtebodavägen 23a, SE-171 65, Stockholm, Sweden; Novo Nordisk Foundation Center for Biosustainability, Technical University of Denmark, DK-2800 Kgs. Lyngby, Denmark; BioInnovation Institute, Ole Måløes Vej 3, DK2200 Copenhagen N, Denmark

**Keywords:** *Saccharomyces cerevisiae*, genome scale metabolic modelling, thermotolerance, Bayesian statistical learning

## Abstract

The molecular basis of how temperature affects cell metabolism has been a long-standing question in biology, where the main obstacles are the lack of high-quality data and methods to associate temperature effects on the function of individual proteins as well as to combine them at a systems level. Here we develop and apply a Bayesian modeling approach to resolve the temperature effects in genome scale metabolic models (GEM). The approach minimizes uncertainties in enzymatic thermal parameters and greatly improves the predictive strength of the GEMs. The resulting temperature constrained yeast GEM uncovered enzymes that limit growth at superoptimal temperatures, and squalene epoxidase (ERG1) was predicted to be the most rate limiting. By replacing this single key enzyme with an ortholog from a thermotolerant yeast strain, we obtained a thermotolerant strain that outgrew the wild type, demonstrating the critical role of sterol metabolism in yeast thermosensitivity. Therefore, apart from identifying thermal determinants of cell metabolism and enabling the design of thermotolerant strains, our Bayesian GEM approach facilitates modelling of complex biological systems in the absence of high-quality data and therefore shows promise for becoming a standard tool for genome scale modeling.

## Introduction

Temperature is the most common environmental and evolutionary factor that shapes the physiology of living cells. Organisms have successfully adapted to survive in diverse temperature ranges^1-3^, where minor deviations from the optimal temperature by merely a few degrees can dramatically impair cell growth. For instance, the model eukaryotic organism *Saccharomyces cerevisiae* has an optimal growth temperature of ~30°C, whereas a temperature of 42°C is already lethal to the organism^4,5^. Since cell growth fundamentally requires all cellular components to be functional in the temperature window of cell growth, proteins, the most abundant group of biomolecules that carry out the majority of catalytic functions and are also the most sensitive to changes in temperature^5-7^, are considered to have the largest effect on cell physiology in relation to temperature. However, despite all our knowledge of temperature effects at both the cellular and molecular levels, including recent breakthroughs in temperature-dependent protein folding^7-10^ and enzyme kinetics^11,12^, the temperature association between proteins and cell physiology is still poorly understood.

Multiple studies have attempted to model the temperature effects on cell growth with very few proteome wide parameters. For instance, the dominant activation barrier and the number of essential proteins to cell growth^13^, activation energy of the growth process and the free energy change of protein denaturation^14^ and others (reviewed in^15^). These models showed excellent performance when describing the general cell growth rate at various temperatures, however, they could not pinpoint the specific rate-limiting enzymes, nor predict the amount of improvement in growth rate by replacing these enzymes with temperature-insensitive homologs.

To this end, genome-scale metabolic models (GEMs)^16-18^, which are a comprehensive mathematical representation of cellular biochemical reactions^19^, have been used to model the thermosensitivity of metabolism in *Escherichia coli*, for instance by associating metabolic reactions with protein structures^20^ or by modelling protein-folding networks^21^. It however remains challenging to model more complex, eukaryotic organisms, such as *S. cerevisiae*, due to their metabolic complexity^16^ as well as due to the lack of availability of the required enzymatic data^7,22^, including high quality protein structures^20,21^. In addition, such GEMs rely on thousands of parameters to describe the temperature effects on protein folding and kinetics^16^, which have to be empirically or computationally estimated^20,21^. This leads to large statistical uncertainties in model parameters and can make the models unreliable, due to inaccurate temperature associations between proteins and cell physiology. Therefore, in order to enable accurate modelling of the temperature dependence of cell metabolism, a key requirement is to develop a modelling approach that resolves the issues with large uncertainties of temperature related parameters and produces accurate temperature constrained predictions.

Hence, in the present study we introduce a Bayesian genome scale modelling approach to model the temperature effect on cellular metabolism in *Saccharomyces cerevisiae*, the most widely used industrial organism with the availability of multiple thermal experimental data^5,23,24^ and highly sophisticated GEMs^16,18,25^. We first quantify and reduce the large uncertainties in the parameters describing enzyme thermosensitivity using Bayesian statistical learning^26^ to simulate phenotypic data. We show that the resulting models are capable of reproducing various experimental datasets and provide explicit insight into how yeast metabolism is affected by temperature. Our approach identifies the sterol metabolism as a key factor in the yeast thermal adaptation, and predicts the flux-controlling enzymes in superoptimal temperature ranges as potential targets for future design of thermotolerant yeast strains. We then experimentally validate the predicted most rate-limiting enzyme by replacing it with an ortholog from a known thermotolerant yeast *Kluyveromyces marxianus.* We hereby demonstrate the power of Bayesian genome scale modelling for studying complex biological systems.

## Results

### GETCool: Using Bayesian statistical learning to integrate temperature dependence in enzyme-constrained GEMs

In this study, we developed a novel approach for incorporating temperature dependence into an enzyme-constrained GEM (ecGEM)^16^ (Fig 1) with the resulting model termed enzyme and temperature constrained GEM (etcGEM). The approach combined the following steps: (i) etcGEM construction (Fig 1a-d), (ii) flux balance analysis (FBA) and (iii) Bayesian statistical learning (Fig 1e). The ecGEM, which includes, besides the traditional stoichiometric matrix, also enzyme abundances and activities, provided an excellent template to directly integrate the enzyme temperature effects. Firstly, for a given reaction, the flux cannot exceed the capability of the enzyme, which is defined as the product of the functional enzyme concentration [*E*]_*N*_ and its *k_cat_*. Secondly, the total amount of enzymes that the cell can afford is also limited^27^. Inclusion of temperature constraints into ecGEM was thus achieved by making [*E*]_*N*_ and *k_cat_* temperature dependent, and by incorporating the additional cost of enzymes in the denatured state (Fig 1a, Method M1). Three thermal parameters were required for each enzyme in the resulting etcGEM, including (i) the melting temperature *T_m_* (Fig 1b), (ii) the heat capacity change 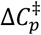(Fig 1c) and (iii) the optimal temperature *T_opt_* (Fig 1d Method M2). Moreover, to capture the temperature effects on the energy cost of non-growth associated maintenance (NGAM), a temperature dependent NGAM expression term was estimated from experimental data and included in the model.

**Fig 1.**
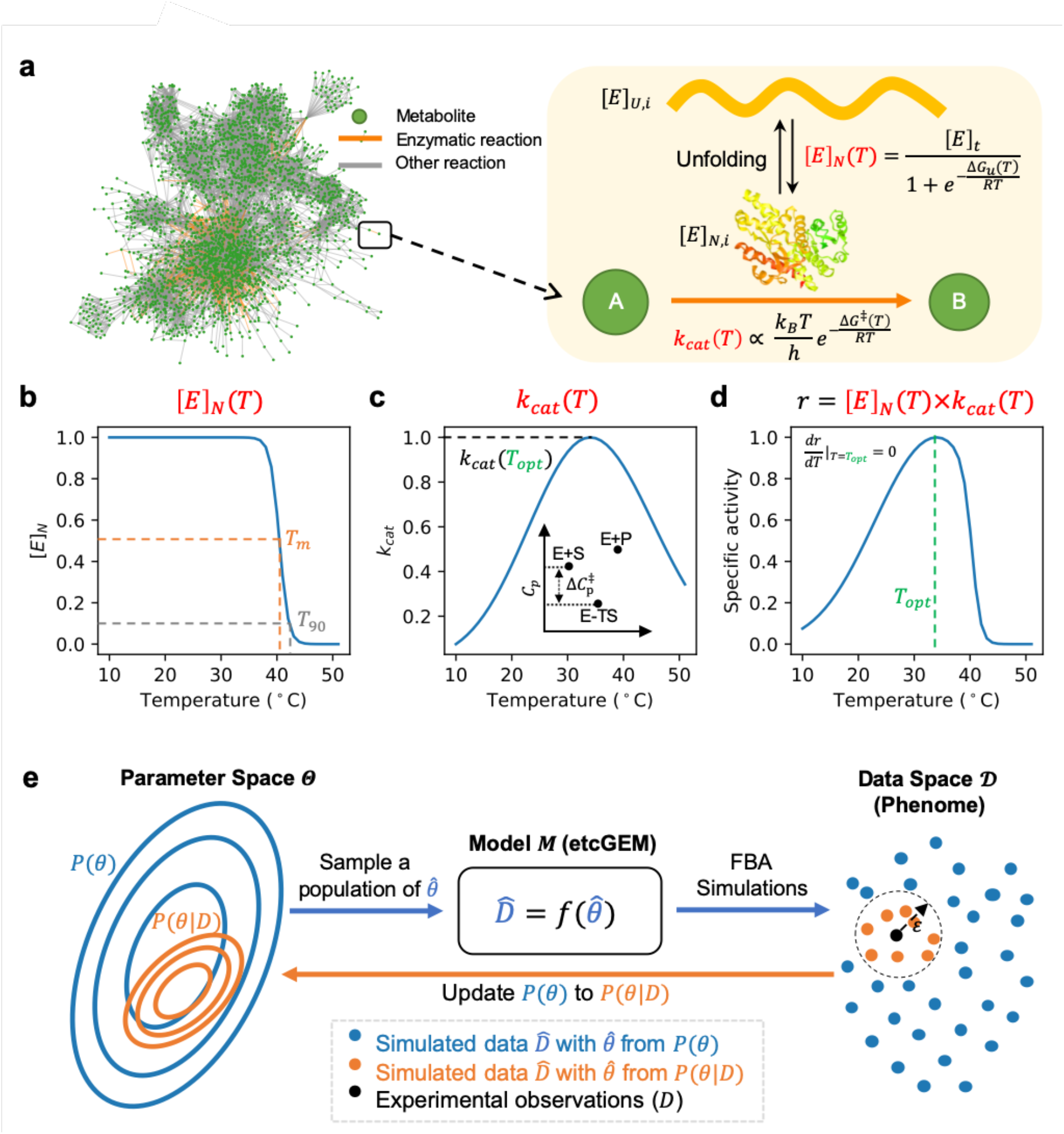
Using Bayesian statistical learning to integrate temperature dependence in enzyme-constrained GEMs. (a) An illustration of the temperature effects on enzyme-catalyzed reactions and their integration into an etcGEM (see detailed description and equations in Methods M3). The metabolic network ecYeast7.6^16^ is shown. (b) A two-state denaturation model^20,21,31^ was used to describe the temperature dependent unfolding process. [*E*]_*N*_ is the concentration of the enzyme in native state; *T*_opt_ is the optimal temperature at which the specific activity is maximized; *T*_m_ and *T*_90_ are temperatures at which there is a 50% and 90% probability that an enzyme is in the denatured state, respectively. (c) Macromolecular rate theory^32,33^ describing the temperature dependence of enzyme *turnover number k_cat_*. Inset shows the heat capacity difference between ground state (E+S) and transition state (E-TS), adapted from Hobbs J., *et al*^32^. (d) Temperature dependence of enzyme *specific activity r*, which is a product of (b) and (c). (e) Overview the Bayesian statistical learning approach, where the problem can be formulated as: given a *generative model (M)* (etcGEM in this study) corresponding to a set of parameters *θ* and a set of measurements *D*(phenome data), Bayes’ theorem provides a direct way of updating the *Prior* distribution of parameters *P*(*θ*) to a *Posterior* distribution 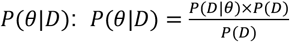. *P*(*θ*|*D*)is thereby a less uncertain description of the real *θ*. Since *P*(*D*|*θ*) is, in most applications, computationally expensive or even infeasible to obtain, an Sequential Monte Carlo based Approximate Bayesian Computation (SMC-ABC)^34^ approach was implemented (Methods M3) to sample a list of parameter sets from the *Posterior*.

To resolve the challenges arising from the uncertainties in the parameter values, we used Bayesian statistical learning^26^, which is a probabilistic framework that has been successfully applied for quantifying and reducing uncertainties in various fields including deep learning^28^, ordinary differential equations^29^ and biochemical kinetic models^30^. The approach uses experimental observations (*D*) to update *Prior* distributions (*P*(*θ*)) of model parameters to *Posterior* ones (*P*(*θ*|*D*)) (Fig 1e). We refer to the model equipped with *θ* sampled from *P*(*θ*) or *P*(*θ*|*D*) as a *Prior* or *Posterior* etcGEM, respectively. The resulting *Posterior* etcGEMs provided a more reliable platform to study the thermal dependence of cell metabolism, with an inherent benefit that the uncertainty in the interpretation and prediction from the improved *Posterior* etcGEMs could also be quantified.

### Bayesian modelling improves etcGEM performance by reducing parameter uncertainties

We next applied the GETCool approach to model the temperature dependence of yeast metabolism. This was done by incorporating temperature effects into the ecYeast7.6^16^ model and the resulting model was termed etcYeast7.6. Enzyme *T_m_* and *T_opt_* parameters were either collected from literature or predicted by machine learning models (Methods M4). The heat capacity change 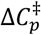 was estimated as −6.3 kJ/mol/K by fitting the macromolecular rate theory to the yeast specific growth rate at various temperatures^32^ and then applied for all enzymes. As a result, the etcYeast model was obtained with an expansion of 2,292 temperature-associated parameters for a total of 764 metabolic enzymes (Fig 1a). The temperature dependence of NGAM was inferred from experimental data (Methods M4, Fig S1).

We observed that etcYeast predictions made using the initial parameter values could not correctly recapitulate experimental observations (Fig S2, Method M5), which included (i) the maximal specific growth rate in aerobic^4^ batch cultivations, (ii) anaerobic^5^ batch cultivations, and (iii) fluxes of carbon dioxide (CO2), ethanol and glucose in chemostat cultivations^23^, at various temperatures. This was however not surprising due to the high level of uncertainty and low accuracy associated with the initial parameter values, as with the experimentally measured *T_m_* we estimated an average standard variance of 3.4 °C, whereas this increased up to 13 °C with the *T_opt_* values predicted by machine learning (Methods M4). For enzymes without experimentally measured *T_m_*, the average of the existing experimental values was used, where the standard variance was 5.9 °C (Methods M4). Another potential source of error was due to assuming the same 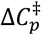 values for all enzymes. We therefore applied the Bayesian statistical learning approach. Here, we first used a three-fold cross validation showing that the above three datasets showed both overlapped and orthogonal information between each other in the Bayesian modelling approach (Fig S3). We then used all three datasets to sample 100 *Posterior* etcGEMs, where each model achieved an average *R*^2^ higher than 0.9 on all three datasets (Fig S4) and could therefore accurately describe the observed measurements (Fig 2a-c and Fig S5). The increased performance on all three datasets clearly demonstrated the need to update the parameter *Prior* distribution to a *Posterior* one.

**Fig 2.**
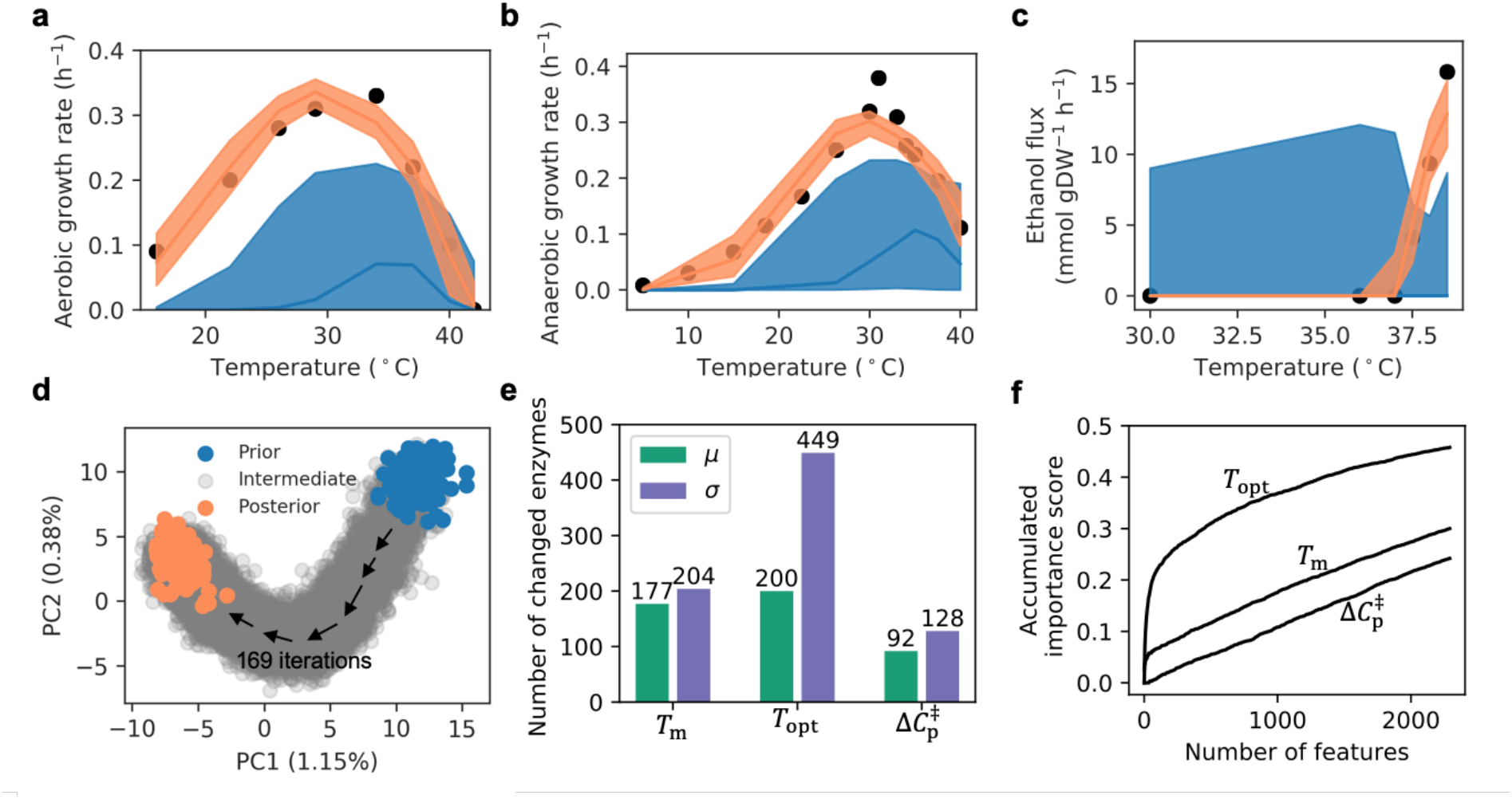
Bayesian modelling improves etcGEM performance by reducing parameter uncertainties. (a-b) Simulated (a) aerobic and (b) anaerobic growth rates in batch cultivations at various temperatures with *Prior* and *Posterior* etcGEMs. (c) Simulated ethanol secretion flux in chemostat at various temperatures. In (a-c), lines indicate median values and shaded areas indicate regions between the 5-th and 95-th percentiles. (d) Principal component analysis (PCA) 21,504 parameter sets 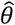 sampled in the Bayesian approach. Each parameter in the set *θ*\* was standardized by subtracting the mean and then be divided by the standard deviation before PCA. 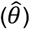 of 128 *Prior* and 100 *Posterior* etcGEMs are highlighted in blue and orange, respectively. All other 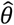 were termed as “intermediate” and marked in grey. (e) The number of enzymes, out of all 764, with a significantly changed mean (Šidák adj. Welch’s *t*-test *p*-value < 0.01) and variance (Šidák adj. one-tailed *F*-test *p*-value < 0.01) in *T_m_, T_opt_* and 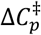 between *Prior* and *Posterior*. Parameters from 128 *Prior* and 100 *Posterior* etcGEMs were used for statistical tests. (f) A random forest model was used to score the importance of all 2,292 parameters during the Bayesian approach (Methods M6). The plot shows the accumulated importance score for each of the three parameter categories.

Next, we explored which parameters had been updated in the Bayesian approach. Principal component analysis of all 21,504 parameter sets generated in the approach showed how the *Priori* distributions were gradually updated to distinct *Posterior* distributions (Fig 2d). Further comparison between *Prior* and *Posterior* distributions revealed that in all three parameter categories, a reduced variance in the updated parameters was more likely than a change in mean values (Fig 2e, protein-wise comparison shown in Fig S6). Particularly for enzyme *T_opt_* s, a significant (Šidák adj. one-tailed *F*-test *p*-value < 0.01) reduction in variance was observed with 59% (449/764), whereas a significant (Šidák adj. Welch’s *t*-test *p*-value < 0.01) change in the mean value was found with merely 26% (200/764). The average standard variance of enzyme *T_opt_*s was thus reduced by almost 50% from ~11 °C to ~6 °C (Fig S6). Importantly, we observed that the approach tended to change the enzyme *T_opt_* rather than its *T_m_* and 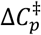 parameters (Fig 2e). In addition, a machine learning approach (Methods M6) further revealed that, out of all three parameter types, the largest contribution to the improved *Posterior* etcGEM performance during the Bayesian approach was from enzyme *T_opt_*s (Fig 2f).

### Yeast growth rate is explained by temperature effects on its enzymes

With the *Posterior* etcGEMs capable of describing various experimental observations (Fig 2a-c), we analysed how the temperature effects on each of the three processes - NGAM, *k_cat_* and the protein denaturation process - contribute to whole cell growth (Fig 3a). We observed that, at temperatures below 29 °C, the temperature dependent *k_cat_* was the only factor that affected the cell growth rate. In the range between 29 and 35 °C, both *k_cat_* and NGAM determined the growth rate. The contribution of enzyme denaturation to the temperature dependence of cell growth, however, was observed only at temperatures higher than 35 °C, with the denaturing effect becoming the dominant effect at ~40 °C and lack of cell growth at 42 °C. Therefore, in contrast to previous reports indicating that an over 10-fold increase in NGAM cost with the temperature change from 30 °C to 33 °C was the major limiting factor to cell growth^5,35^, our modelling approach showed that the increased NGAM has a merely moderate effect on growth rate (Fig 3a).

**Fig 3.**
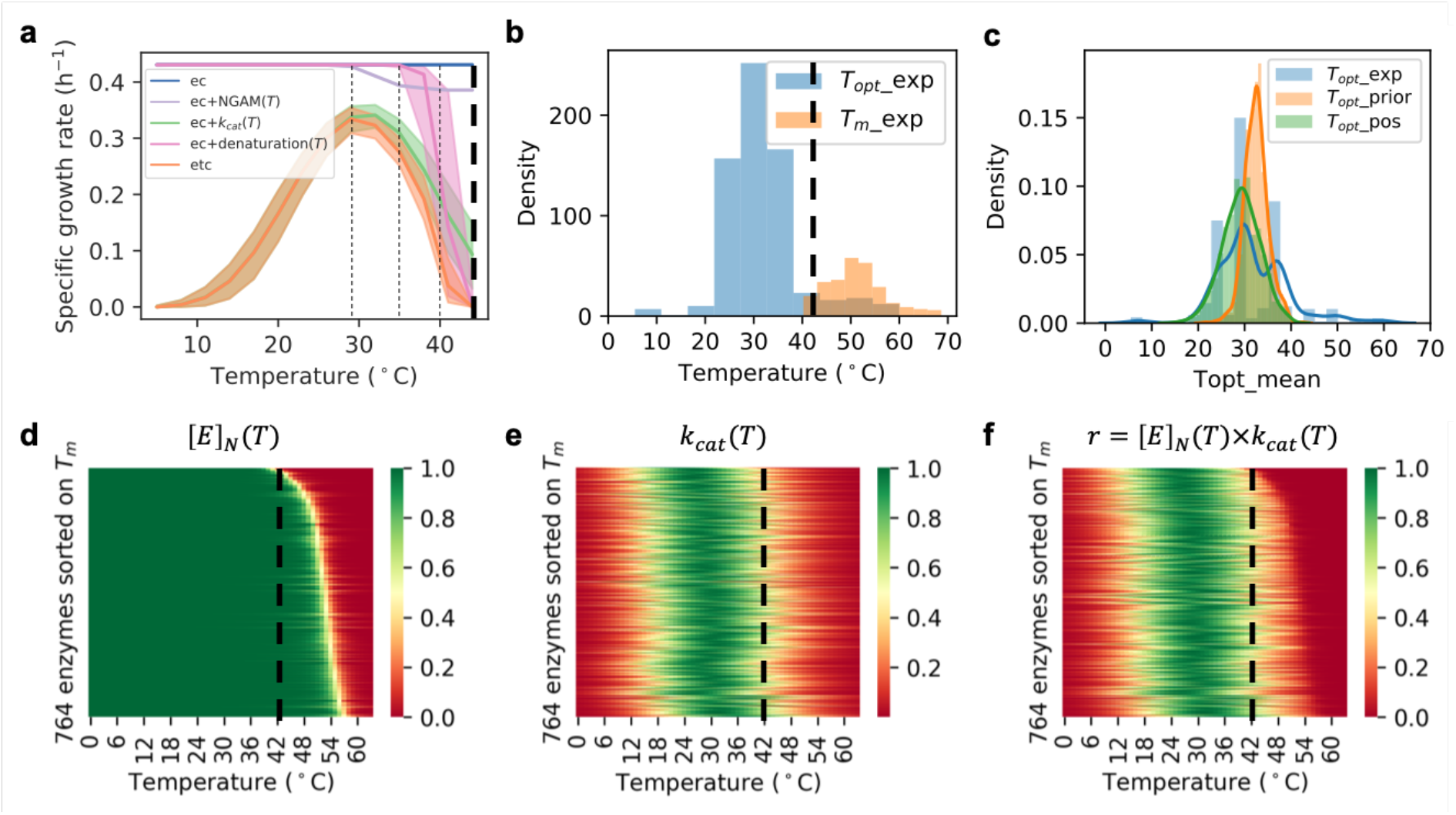
Yeast growth rate is explained by temperature effects on its enzymes. (a) Illustration of how the temperature dependence of different processes combines to affect the growth rate. Fig legend: ec - predictions with the enzyme constrained model; ec+NGAM(T) - incorporates the temperature effects on non-growth associated maintenance into the ec model (Fig SX); ec+kcat(T) - incorporates the temperature effects on enzyme *k_cat_* values into the ec model; ec+denaturation - incorporates the temperature effects on enzyme denaturation into the ec model; etc - enzyme and temperature constrained model that includes the temperature effects on NGAM, *kcat* and enzyme denaturation into ec model. The growth rate at each temperature point was simulated with all 100 posterior etcGEMs. Lines indicate median values and shaded areas indicate regions between the 5th and 95th percentiles. (b) Comparison between distributions of experimentally measured enzyme *T_opt_*s from BRENDA^37^ and *T_m_*s from Leuenberger P *et al*.^7^ in *S. cerevisiae*. (c) Comparison among distributions of mean of *Prior T_opt_*s which were predicted by Tome^22^, mean of Posterior *T_opt_*s and experimental *T_opt_*s from (b). (d) Probability of 764 enzymes in the native state. From top to bottom, the enzymes showed increased *T_m_*s. Each pixel represents one probability value of an enzyme at a specific temperature. (e) Normalized *k_cat_* values of 764 enzymes at different temperatures. Each pixel represents one normalized *k_cat_* value of an enzyme at a specific temperature. (f) Normalized specific activities of 764 enzymes at different temperatures. The values in (f) are products of (d) and (e). In (d,e,f), an equal ordering of enzymes is shown.

Interestingly, the temperature dependence of enzyme *k_cat_*s alone could explain the temperature dependence of cell growth below 35 °C, including the decline in cell growth right after the optimal growth point defined by OGT. According to the macromolecular rate theory^32,33^, *k_cat_* degeneration at temperatures above the optimal point can be attributed to the negative values of 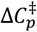 for enzyme catalysis. This can explain the negative curvature of enzyme *specific activities* in the absence of the denaturation process^32,33,36^. Given that experimentally measured enzyme melting temperatures (7%) are on average 20 °C higher than enzyme *T_opt_*s collected from BRENDA^37^ (Fig 3b), protein denaturation alone seems to be insufficient to explain the thermal mechanism underlying enzyme *T_opt_*s. In addition, all posterior *T_opt_*s showed a similar distribution as experimental *T_opt_*s, even though the etcGEM had never seen those experimental *T_opt_*s (Fig 3c), which supported our use of the macromolecular rate theory in the model. This indicates that *k_cat_* degeneration, in addition to protein denaturation, plays an important role in the temperature dependence of yeast cell growth.

We further observed that, even though the model contained only 764 enzymes from a total of ~6,700 proteins^38^, protein denaturation alone could still explain termination of cell growth at 42 °C (Fig 3a). However, in the *Posterior* etcGEMs, only 9 enzymes (1%) with a mean melting temperature below 42 °C were present (ERG1, ATP1, ALA1, KRS1, SER1, HEM1, PDB1, ADH1 and TRP3) (Fig S7), of which three (ATP1, HEM1, PDB1) are located in the mitochondria^39^. The other enzymes remained in the native state even at temperatures several degrees higher than 42°C (Fig 3d), though they were enzymatically active only in the temperature window of cell growth between 10 °C and 42 °C (Fig 3f), due to the low *k_cat_* values beyond this temperature range (Fig 3e).

### Metabolic shifts are explained by temperature-induced proteome constraints

Published reports show that at temperatures above 37°C in chemostat cultures with a dilution rate of 0.1 h^−1^, yeast shifts its metabolism from a completely respiratory one to a partly fermentative one, which is also accompanied by a large increase in glycolytic flux^23^. Since our updated *Posterior* etcGEMs are able to simulate this metabolic shift (Fig 2c and Fig S5), we used them to further explore the mechanisms behind the observed process. We observed that the shift occurs due to a proteome constraint, meaning that the total protein level in the cell reaches an upper bound (Fig 4). The proteome constraint occurs due to the decrease in enzyme specific activities with increasing temperature (Fig 3f) and since the maximal protein amount in the cell is limited^27^. As a result, the cell has to synthesize more enzymes to maintain cell growth at the given growth rate (Fig 4) until the enzyme amount hits the upper bound. This is also consistent with earlier studies showing that the activation of the Crabtree effect in chemostat cultures at 30°C is due to a proteome constraint^16,40^. When the temperature increases above 36 °C, ATP production by glycolysis is dramatically increased, while ATP production by the mitochondria decreases (Fig 4). Even though the respiratory pathway produces more ATP per glucose amount, the fermentative pathway produces more ATP per protein mass and therefore becomes more energetically efficient when the cell reaches a proteome constraint^40^. In addition, three key mitochondrial enzymes (ATP1, HEM1 and PDB1) (Fig S7) were found to be unstable, which make the respiratory pathway even more resource-inefficient for ATP production.

**Fig 4.**
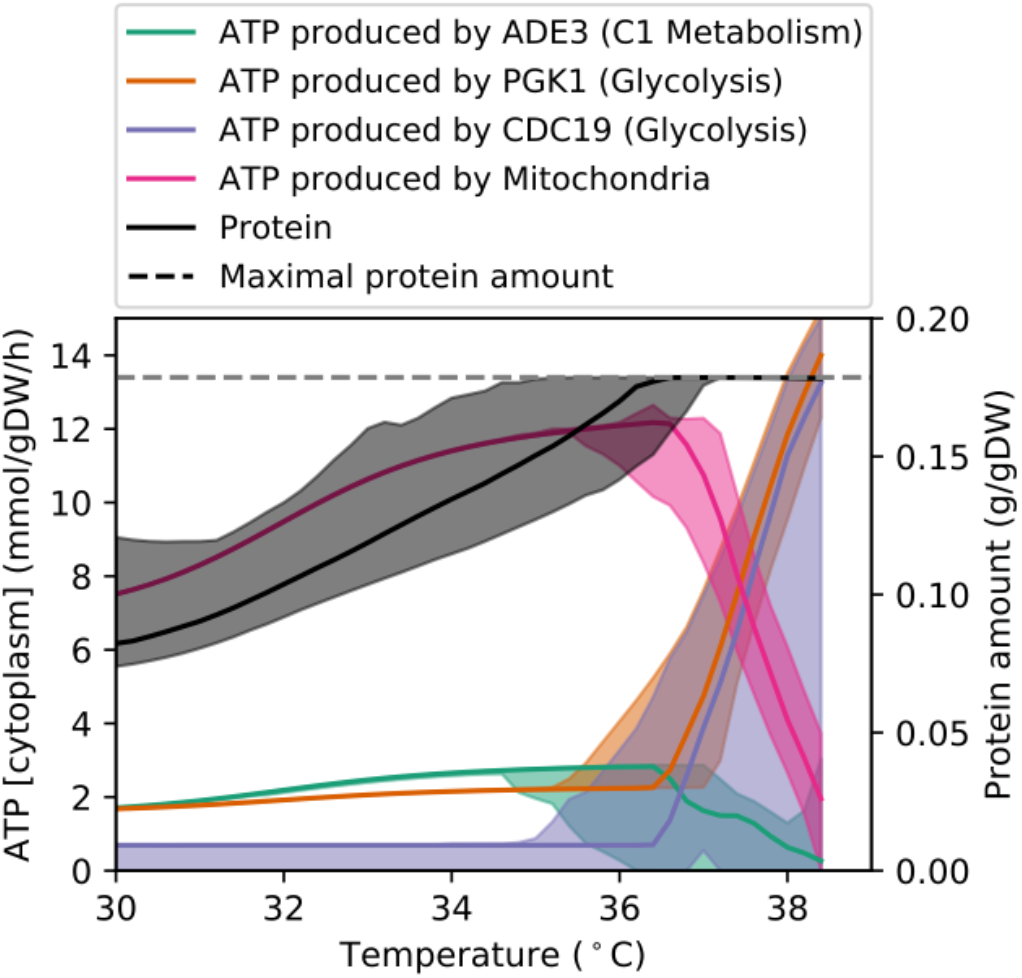
Metabolic shifts are explained by temperature-induced proteome constraints. The ATP production in cytoplasm and the total protein amount required at different temperatures were simulated using *Posterior* etcGEMs with chemostat culture settings with a dilution rate of 0.1 h^−1^ (Methods M4). Lines indicate median values and shaded areas indicate the region between the 5th and 95th percentile.

### etcGEM uncovers growth rate-limiting enzymes

To investigate which enzymes limit the cell growth at superoptimal temperatures, the flux sensitivity coefficient of each enzyme was calculated (Methods M5). Among all the enzymes in the model, the squalene epoxidase ERG1 displayed an order of magnitude higher median flux sensitivity coefficient than other enzymes, indicating that it is the most flux-controlling enzyme at 40 °C (Fig 5a) and above (Fig S8). Furthermore, removal of the temperature constraint on ERG1 increased the simulated specific growth rate from 0.09 to 0.14 h^−1^ (Fig 5b). We therefore evaluated the impact of replacing the wild-type *ERG1* gene with *ERG1* from the thermotolerant yeast *Kluyveromyces marxianus* (kmERG1, Methods M7). At first, at the lethal temperature of 42 °C, only a small improvement in growth rate (from 0.01 to 0.06 h^−1^) was predicted and no significant growth difference was detected between the wildtype and the strain with kmERG1 (Fig S9). However, already after 2 generations of adaptation at 40 °C, the strain with KmERG1 indeed showed significantly better growth than the wild type (Fig 5c).

**Fig 5.**
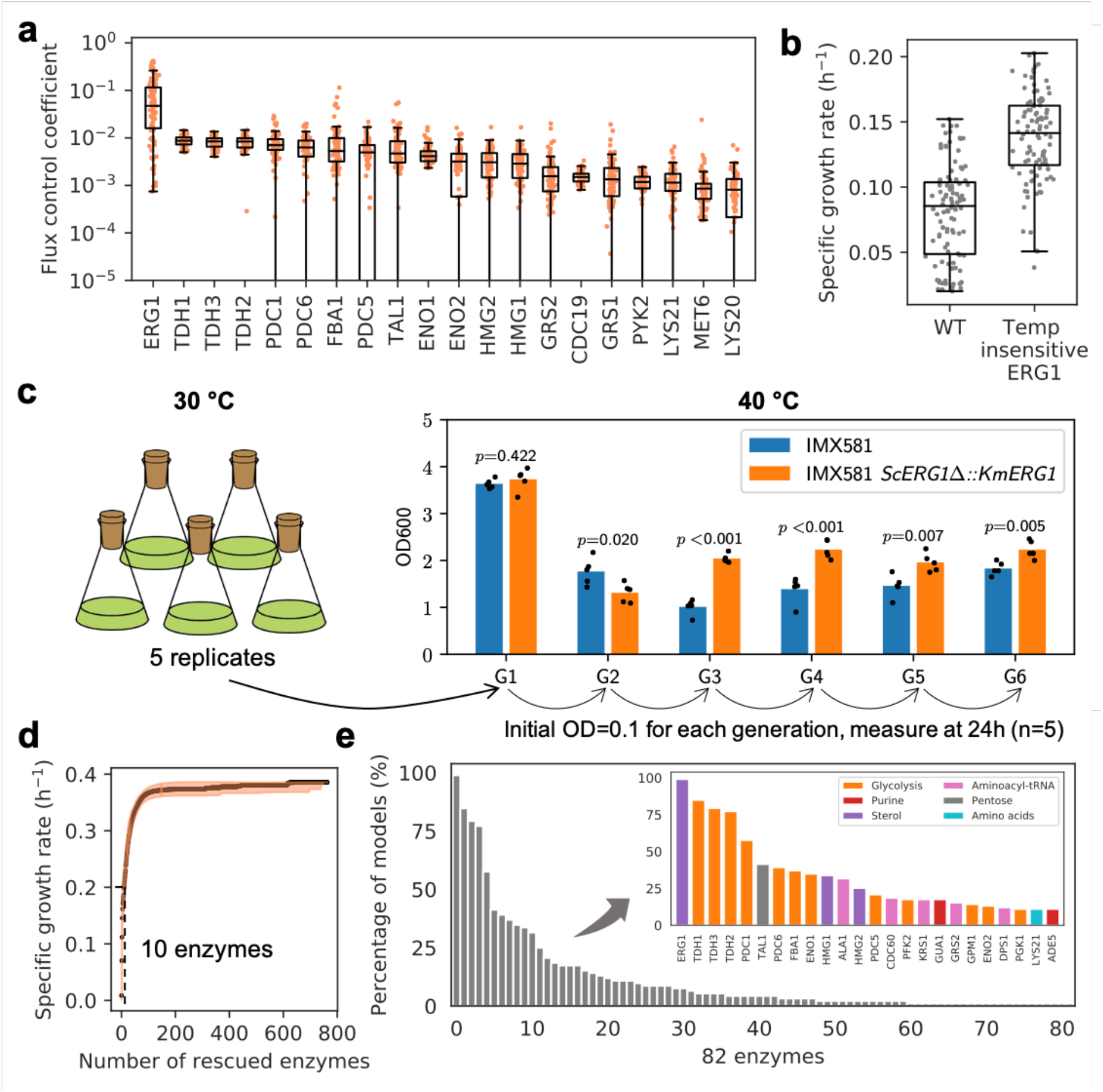
etcGEM uncovers growth rate-limiting enzymes. (a) 20 enzymes with the highest flux sensitivity coefficients at 40 °C. Each dot represents the prediction from one *Posterior* etcGEMs. (b) Predicted maximal specific growth rate of wide-type yeast and the one without any temperature constraints (fully functional) on ERG1 enzyme at 40 °C. (c) The effect of KmERG1 expression on thermo tolerance in *S. cerevisiae*. The strains were cultivated at 40 °C for six generations to reach the steady state of growth. Optical densities (600 nm) are shown at 24 h. Each bar indicates the mean and dots represent the values of 5 replicates. *p*-values denote Welch’s *t*-test. (d) Simulated maximum specific growth rate by removing the temperature constraints of most rate-limiting enzymes at each step in each *Posterior* etcGEM at 42 °C. Lines indicate median values and shaded areas indicate the region between the 5th and 95th percentiles. (e) The percentage of *Posterior* etcGEMs predicts an enzyme to be in the minimal enzyme set required to be fully functional at 42 °C in order to achieve a maximal specific growth rate of 0.2 h^−1^. Inset shows the names and pathways of genes predicted by more than *Posterior* etcGEMs 10% they are involved in.

The reduced growth rate at 42 °C is likely caused by an impaired function of several different enzymes, and rescuing a single enzyme is insufficient to improve the growth rate. Therefore, in order to characterize the set of growth rate-limiting enzymes at 42 °C, we gradually removed the temperature constraints on enzymes (set *k_cat_* and denaturation temperature independent) in the order of decrescent flux sensitivity coefficient values in each of the *Posterior* etcGEMs. Interestingly, in the case of recovering the cell growth rate to 0.2 h^−1^, we found an agreement among all *Posterior* etcGEMs that 10 enzymes are required to be fully functional at 42 °C (Fig 5d). Since each model predicted a different subset of such enzymes, an ensemble approach was used to count the number of models (votes) in which an enzyme is predicted to be one of 10 such enzymes (Fig 5e). In total, 82 enzymes were predicted by at least one *Posterior* etcGEM, and only 24 (out of 82) enzymes were each predicted by more than 10% of the *Posterior* etcGEMs (Fig 5e, inset). Among these 24 enzymes 12 enzymes were engaged with Glycolysis and 3 enzymes were involved in sterol biosynthesis: ERG1, and HMG1,2 catalyzing the flux-controlling steps in sterol biosynthesis^41^. The remaining enzymes were mainly involved in DNA or protein synthesis related pathways.

## Discussion

Here, we present a Bayesian genome scale modelling approach to resolve the temperature dependence of cellular metabolism, termed GETCool. Using an enzyme-constrained GEM^16^ as a template, we modelled the temperature effects on each individual enzyme by including temperature dependent terms for the independent processes of denaturation as well as catalysis (Fig 1A). Due to the high level of uncertainty and low accuracy associated with the initial thermal parameter values (Fig S6), which were a result of experimentally measured noise or variability arising from machine learning or theoretical predictions (Methods M5), the model predictions initially could not correctly recapitulate experimental observations (Fig 2a-c and Fig S2). We therefore used Bayesian statistical learning that enabled updating our *Prior* guess of the highly uncertain thermal parameters to a more accurate *Posterior* estimation of these parameters according to observed phenotypic data (Fig 1e). The resulting *Posterior* etcGEMs accurately describe the experimental observations (Fig 2a-c) and thus provide a more reliable platform to study the thermal dependence of yeast metabolism.

Previous studies modelling the temperature dependence of enzyme activities have relied mainly on protein denaturation and the Arrhenius equation, where protein denaturation explained the negative curvature for temperature dependence of enzyme activity^20,21^. However, with the increasing amount of evidence showing that protein denaturation alone is insufficient to explain the decrease in enzyme specific activity above *T*opt, macromolecular molecular rate theory^32,36^ has become a promising alternative. It was successfully applied to many enzymes^32,33,36^, including its use in explaining the evolution of enzyme catalysis^36^. According to the theory, a negative heat-capacity change 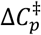 exists between the transition state and the ground state in the enzyme catalytic process (Fig 1c), which leads to a negative curvature for temperature dependence of enzyme activity in the absence of denaturation^32^. We found that with this theory, temperature dependence of *k*_cat_s acts as a major contributor to the cell growth rate at all temperatures, which can especially explain the decline in cell growth rate right after the optimal growth temperature (Fig 3a). Yeast enzymes only maintain high *k*_cat_s in the temperature window of cell growth (Fig 3e), which means that the metabolism becomes inefficient at superoptimal temperatures due to the general decrease in enzyme turnover without denaturations (Fig 3d-f).

Using the Bayesian genome scale modelling approach to quantitatively depict the temperature effects on yeast metabolism led to insights into the long-standing discussion on the roles of different cellular factors in cellular fitness under heat stresses^4,5,7,23,42^. For instance, protein denaturation has been suspected as one of the main causes of the decline in cell growth beyond the optimal growth temperature point. However, recent high throughput measurements of melting temperatures (*T*_m_) for 707 *S. cerevisiae* proteins revealed a *T*_m_ distribution with a mean value of 52 °C and a minimum of 40 °C^7^, which suggests that protein denaturation alone might not be sufficient to explain the decline of yeast cell growth between 30°C (optimal growth temperature, OGT) and 42°C (lethal temperature point). An alternative explanation is provided by the evidence of a significant increase of non-growth associated ATP maintenance (NGAM) observed with yeast cells grown in anaerobic chemostat cultivations at high temperatures (33-40°C) compared to ones grown at low temperatures (5-31 °C)^5^, which suggests an imbalance in cellular energy allocation in the superoptimal temperature range. Quantitative assessment using our modelling approach revealed that the impaired cell growth is caused by a combination of decreased *k*_cat_ values, increased NGAM costs and protein denaturation (Fig 3). Furthermore, between 30 and 35 °C, the combined decrease in *k*_cat_s and increase in NGAM explains the decline in cell growth, whereas with temperatures above 35 °C, protein denaturation becomes the dominant factor, causing cell death at 42°C. However, in accordance with published findings that cellular proteomes have a broad distribution of protein stability with only proteins at the tail of the distribution being problematic^43^, using our approach we identified only ~1% unstable enzymes denatured at the lethal point (*T*_m_ lower than 42 °C, Fig 3d).

We identified two interesting metabolic pathways involved in yeast thermotolerance: sterol metabolism and mitochondrial energy metabolism. With sterol metabolism (Fig 5d), it is known that high sterol levels help yeast cells survive under heat stress^44^ and changes of the sterol composition of the yeast membrane from ergosterol to fecosterol^45^ can significantly increase yeast thermotolerance. However, yeast was found to downregulate its whole ergosterol biosynthesis at both transcription and translation levels when increasing the temperature from 30°C to 36 °C (Fig S10). Our modeling approach identified three problematic enzymes (Fig 5d: HMG1,2 and ERG1) in the sterol metabolism, which are also flux-controlling enzymes in the sterol biosynthesis pathway^46^. We experimentally confirmed that replacement of ERG1 with its ortholog in the thermotolerant yeast *K. marxianus* can significantly improve the cell growth at 40 °C (Fig 5c). We thereby hypothesize that, since those three enzymes are problematic at superoptimal temperatures, there is no need for the cell to maintain high expression and translation levels of other enzymes in the same pathway. Instead, it has to downregulate its whole ergosterol biosynthesis to save resources and increase fitness.

With mitochondria, previous studies have indicated that the mitochondrial genome plays an important role in yeast thermal adaptation^47–49^. We found that out of the 9 unstable enzymes identified with the *Posterior* etcGEMs (with a *T*_m_ lower than 42 °C, Fig S7), three (ATP1, HEM1 and PDB1) belonged to the mitochondrial energy metabolism. Simulation of chemostat data (Fig 4) revealed that at superoptimal temperatures, yeast prefers to produce ATP via the glycolysis metabolism instead of the mitochondrial energy metabolism in the mitochondria. Furthermore, mitochondria only exists in eukaryotes and almost all of them have evolved to have an optimal growth temperature below 40 °C^3^. All these findings indicate that mitochondria are not evolved to be functional at very high temperatures (e.g. >42 °C). Since mitochondrial energy metabolism is not essential for yeast cell growth, as there are alternative energy pathways (Fig 4), this also explains why we could not successfully predict mitochondrial enzymes to be engineering targets for the recovery of cell growth at 42 °C (Fig 5d), despite the existence of three unstable enzymes in the mitochondrial energy metabolism.

In conclusion, we demonstrate the usefulness of a Bayesian genome scale modeling approach for reconciling temperature dependence of yeast metabolism. Describing the link between temperature and cell physiology is of industrial importance, e.g. for finding optimized production of biochemicals^24,50–52^, but also in medicine, e.g. to understand the effects of temperature on human metabolism^53–55^. Furthermore, based on its success here, we foresee that our method can be integrated into genome scale modelling approaches in general. This approach can also become a staple of GEM modeling in order to resolve uncertainties present in the data, which can be important as GEMs have become a widely used platform for integration of various biological data, such as transcriptomics and proteomics data that are associated with large uncertainties^56^.

## Materials and Methods

### M1. A temperature dependent enzyme-constrained genome scale metabolic model (etcGEM)

The central concepts of an enzyme constrained model^16^ are: 1) the flux through each reaction cannot exceed the capacity of its catalytic enzyme: *υ_i_* ≤ *k_cat,i_* · [*E*]_*i*_, where [*E*]_*i*_ is the concentration of enzyme *i*; 2) the total enzyme amount is constrained by the experimental measurement: ∑ [*E*]_*i*_ < [*E*]_*t*_. Once the temperature dependent denaturation and *k_cat_* were considered, [*E*]_*i*_ in the first constraint should be [*E*]_*N,i*_ which is the concentration of individual active enzymes. [*E*]_*i*_ in the second constraint should be [*E*]_*t,i*_ = [*E*]_*N,i*_ + [*E*]_*U,i*_, which is the total concentration of enzymes in both active and denatured forms (Fig 1a). In addition, to capture the increased expenditure for maintenance under increased heat stress, a temperature dependent Non-Growth Associated ATP maintenance term can be assumed from experimental measurements. In summary, the updated constraints constraints in etcGEM are

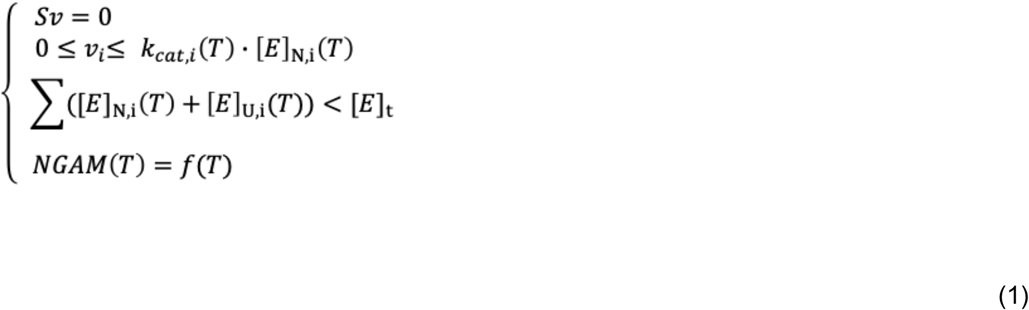

The effect of temperature on *k_cat_* values can be described with an expanded Arrhenius equation (macromolecular rate theory), by including a non-zero heat-capacity change 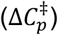 between the transition state and the ground state of the enzyme catalytic process^32,33^:

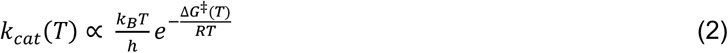

in which *k*_B_ is the Boltzmann constant, *h* is Planck’s constant, *R* is the universal gas constant, and Δ*G*^‡^(*T*) is the free energy difference between the ground state and the transition state. The latter can be expanded as

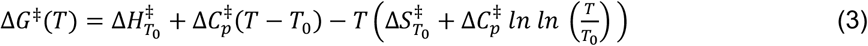

where 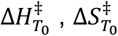 and 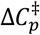 are the differences in enthalpy, entropy and heat capacity change between the transition and ground states, respectively, and *T*_0_ is the reference temperature. This theory has been successfully applied to study the temperature dependence of enzyme activity^32,33^ and evolution^36^.

Since there is not enough detailed information regarding the heat-induced denaturation process of yeast proteins, a simple two-state model denaturation was assumed as in many other studies^20,21,31^. In such a model, a protein molecule could be either in a native state (N) or a denatured state (U), and an equilibrium state was assumed: *N ↔ U*. Thereby

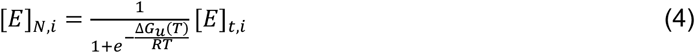

in which [*E*]_*t,i*_ = [*E*]_*N,i*_ + [*E*]_*t,i*_, where [*E*]_*t,i*_is the concentration of enzyme *i* and Δ*G_U_*(*T*) is the free energy difference between the denatured state and the native state and can be expressed as

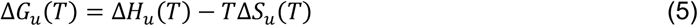

where Δ*H_u_*(*T*) and Δ*S_u_*(*T*)are the enthalpy and entropy changes between the denatured and native states at temperature *T*. It has been found that convergence temperatures 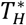 (373.5 K) and 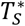 (385 K) exist for Δ*H_u_* and Δ*S_u_*respectively^42,57,58^. At such temperatures, the Δ*H_u_* and Δ*S_u_* converge to a common value of Δ*H*\* and Δ*S*\*. Thereby,

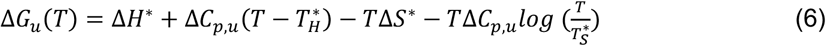

in which Δ*C_p,u_* is the difference in heat-capacity change between the denatured and native states.

In summary, the values of Δ*G*^‡^(*T*) and Δ*G_u_*(*T*) need to be determined in order to model the temperature dependence of enzyme activities, and they can be associated with six unknown parameters: 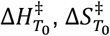 and 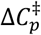 for Δ*G*^‡^(*T*), and Δ*H*\*, Δ*S*\*and Δ*C_p,u_* for Δ*G_u_*(*T*).

### M2. Computation of thermal parameters

Since it is difficult to directly measure those six thermal parameters (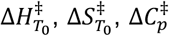, Δ*H*\*, Δ*S*\*and Δ*C_p,u_*) for each enzyme, indirect measurements have to be used to approximate the larger set of thermal parameters. As there are six free variables in the system, six different equations are required to solve for those parameters.

1. At the protein melting temperature *T_m_*:

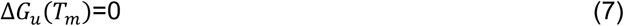
2. At the enzyme optimal temperature *T_opt_*, the enzyme activity is maximized:

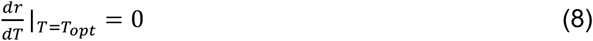

in which *r = k_cat_*[*E*]_*N*_;
3. *k_cat_* at the enzyme optimal temperature *T_opt_* is known:

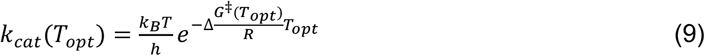
4. 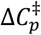 value can be approximate from temperature dependence of cell growth rate^32^
5. We found that there is a very strong linear correlation (*r*^2^ = 0.998, *Pearson*’s correlation) between Δ*H*\*and Δ*S*\* of 116 proteins from Sawle L et al^42^ (Fig S9)

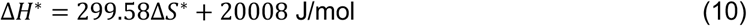
6. For some enzymes, *T*_90_, where a 90% possibility exists that an enzyme molecule is in the denatured state, is experimentally measured:

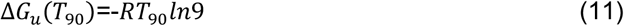

As a result, the six thermal parameters 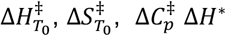, Δ*S*\*and Δ*C_p,u_* can be obtained by solving the above equations.

In the case of lacking *T*_90_ or failed to obtain a positive Δ*C_p,u_*, protein sequence length was used to estimate Δ*H*\* and Δ*S*\*^42^ as below:

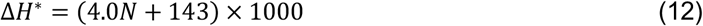

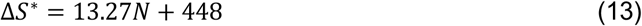

### M3. Sequential Monte Carlo based Approximate Bayesian Computation (SMC-ABC)

Approximate Bayesian Computation^34^ was applied to infer parameter sets from *Posterior* distributions. Given an observed dataset *D* and a model specified by 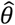 sampled from the *Prior* distribution *P*(*θ*), if the distance between simulated data 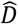 and observed *D* is less than a given threshold *ϵ*, then this 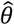 is accepted as the one sampled from 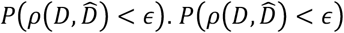 is often used to approximate the *Posterior P*(*θ|D*) when *ϵ* is sufficiently small. In case of high-dimensional parameter space and/or when the *P*(*θ*) is very different from *P*(*θ|D*), the acceptance rate would be very low and thus this approach becomes computationally expensive to generate a population of 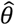 from 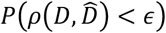. In this work, a sequential Monte Carlo approach was designed as follows to generate a population of 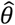 sampled from 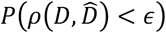:

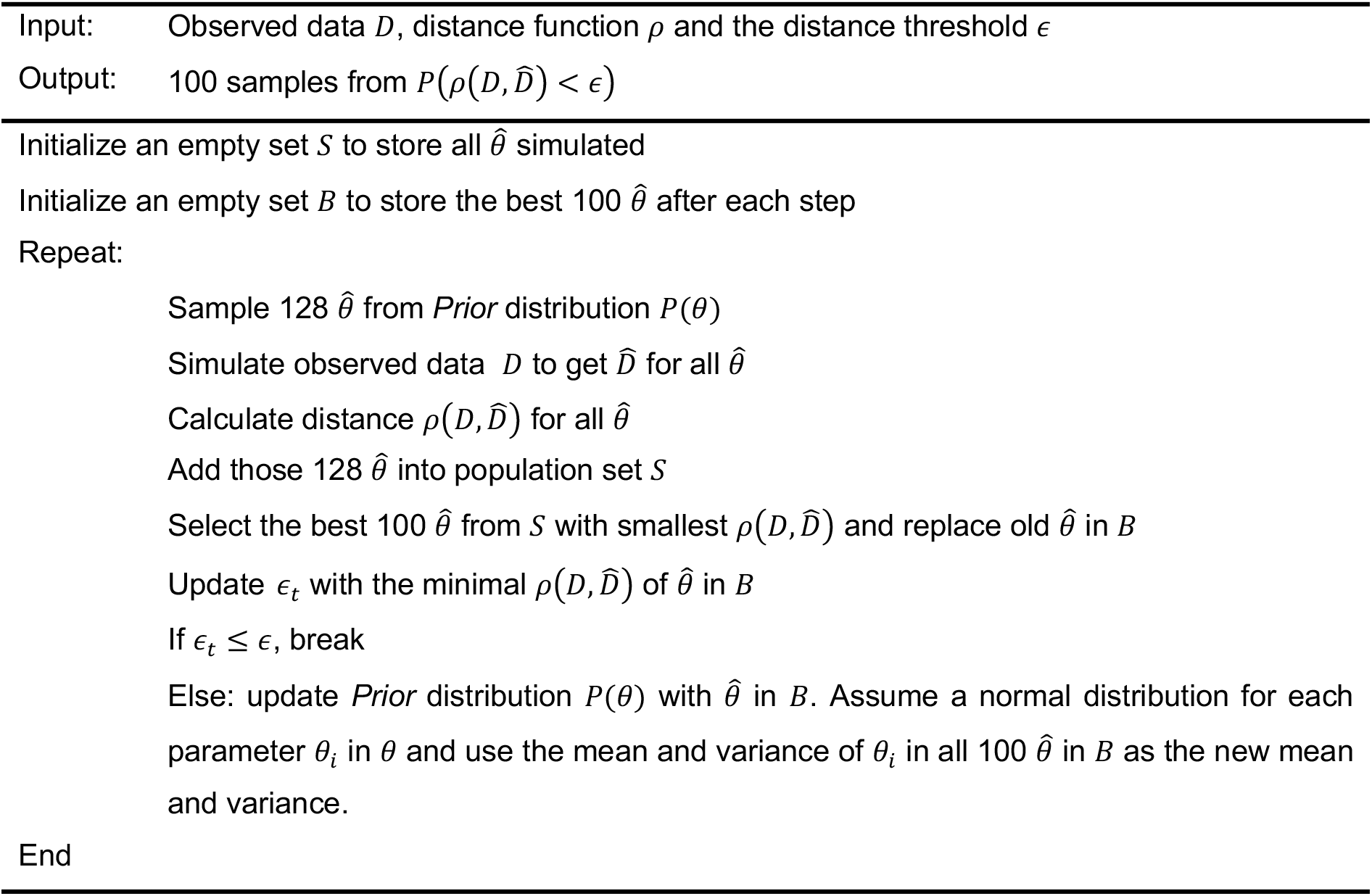

### M4. Collection and estimation of enzyme thermal parameters in etcYeast7.6

The enzyme-constrained model for yeast with minimal medium was taken from^16^.

#### Melting temperatures

Among the 764 enzymes included in ecYeast7, the *T_m_* (melting temperature) and *T*_90_ (the temperature at which 90% of the protein is in the denatured state) for 266 yeast proteins have been reported previously^7^. For enzymes lacking an experimentally measured *T_m_*, a melting temperature of 51.9 °C (the average of existing *T_m_*s of 707 yeast proteins) was assumed. In the original paper^7^, the 95% confidence interval was reported for peptides measured in the experiments and the average standard error was estimated at 3.4 °C. This same value was used as the uncertainty measure for the experimentally determined *T_m_*s, since the standard error for protein *T_m_* was not available. The *T_m_* of the 266 enzymes was then described with a normal distribution *N*(*T_m,i_*,3.4), in which *T_m,i_* is the experimentally measured melting temperature of protein *i*. For enzymes that uses the mean *T_m_* of 707 proteins^7^ as *T_m_* estimation, the corresponding uncertainty is described as the the standard deviation of the the 707 *T_m_*s, equalling 5.9 °C. Thereby, a normal distribution *N*(51.9,5.9) was used.

#### Enzyme optimal temperature

*T_opt_* values of all enzymes in this study were calculated using a previously described machine learning method^22^, which predicts enzyme *T_opt_*based on primary sequences. This model has a coefficient of determination (R2 score) of 0.5 on the test dataset. Root-mean-squared error (RMSE) of the prediction was then estimated with:

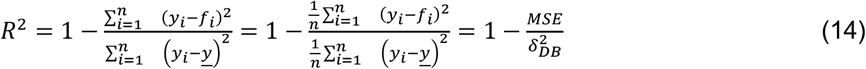

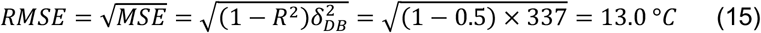

where *f_i_* is the predicted value and *y_i_* is the observed true value of enzyme *i*. Then each one of these predicted *T_opt_*s was described with a normal distribution *N*(*T_opt,i_*, 13.0).

#### Heat capacity change

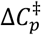 value was approximated by assuming temperature dependence of yeast cell growth rate as −6.3 kJ/mol/K for all enzymes^32^. Given that 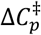 should be in general negative for most enzymes^33^, a standard variance of 2.0 was selected from testing a wide range values because it covers a broad range of 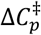 and with a very low possibility of getting a positive value (Fig S10). A normal distribution of *N* (−6.3,2.0) was subsequently used to describe the 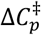 of all enzymes.

#### NGAM

To capture the increased expenditure for maintenance under increased heat stress, an empirical equation (Fig S1) was constructed to estimate the Non-Growth Associated ATP Maintenance at different temperatures:

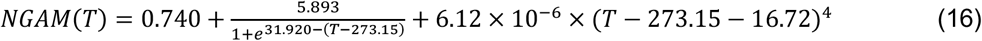

based on the experimental data^5^. Since the experimental data only covers the temperature range of between 5-40°C, any NGAM for temperatures lower than 5°C was set to the value at 5°C and for those higher than 40°C was set to the value at 40°C. The equation (16) was used for the anaerobic growth data as well as for aerobic growth, since no experimental data was available for this condition.

### M5. FBA simulations with etcYeast7.6

At a given temperature, first the *k_cat_* values and 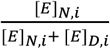 were calculated and integrated into the enzyme-constrained model and then the NGAM at this temperature was calculated and included in the model.

#### Batch cultivation

For batch growth simulations, unlimited substrates were used, the same as described in^16^. The enzyme saturation factor *σ* of 0.5 was used^40^. For simulation of anaerobic growth, in addition to the above changes, the uptake of oxygen was blocked and fatty acids and sterols were supplied into the medium as described in^16^. The growth associated ATP maintenance (GAM) was estimated from experimental data^5^ as 70.17 mmol ATP/gdw. Other parameters were unchanged.

#### Chemostat cultivation

For the simulation of fluxes at aerobic chemostat conditions, with the same model settings as aerobic batch condition, the simulation was carried out by first fixing the growth rate to a given dilution rate (0.1 h^−1^) and minimizing the glucose uptake rate. Then the glucose uptake rate was fixed to the simulated value multiplied by a factor of 1.001 (for simulation purposes). Finally, the total enzyme usage was minimized (same as used in^16^).

#### Flux Sensitivity Analysis

To get the flux sensitivity coefficient of an enzyme at a given temperature, the *k_cat_* of all reactions that associated with this enzyme were perturbed by a factor of (1 + *δ*). Then the maximal growth rates were simulated before (*u*) and after (*u_p_*) perturbation. Finally, the flux sensitivity coefficient of enzyme *i* was calculated as 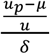, where *μ* and *u_p_* are maximal specific growth rate before and after perturbation. *δ* of 10 was used in this study.

### M6. Analysis of models generated with the Bayesian approach

#### Distance function

The observed data used in this study was the maximal specific growth rate in aerobic^4^ and anaerobic^5^ batch cultivations at different temperatures, and glucose, carbon dioxide and ethanol flux values at different temperatures measured in chemostat cultivations with a dilution rate of 0.1 h^−1 23^. The distance function was designed as follows: first, the coefficient of determination (*R*^2^) between simulated and experimental data was calculated for each of the above conditions. Then the average *R*^2^ across these three conditions multiplied by −1 was used to represent the distance 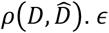 of −0.9 was used in the SMC-ABC simulation.

#### Statistical tests for comparison between *P*(*θ*) and *P*(*θ*|*D*)

The significance test for the difference in mean values between *Prior* and *Posterior* was carried out by Welch’s t-test^59^. The significance test for reduced variance was carried out by the one-tailed *F*-test. *p*-values were adjusted with the correction^60^ using a family-wise error rate of 0.01. The significance cutoff was set to 0.01 (Fig 2e).

#### Machine learning applied to score the importance of parameters

2292 parameters of 21,504 parameter sets were used as the input feature matrix and the average *R*^2^ scores obtained with the Bayesian approach were used as target labels. The dataset was split into train (80%), validation (10%) and test (10%) datasets. A random forest regressor with 1000 estimators was used. The train and validation datasets were used to optimize the hyper-parameter. The obtained model could explain in total 23% the variance in the test dataset. The feature importance scores were extracted directly from the obtained model.

### M7. Experimentally validate ERG1

#### Genetic Manipulation

The background strain we used in this study was IMX581 derived from CEN.PK113-5D, which contains an integrated Cas9 expression cassette controlled by TEFp promoter^61^. All the genetic manipulations were conducted based on the CRISPR/cas9 system. The codon-optimized kmERG1 were ordered from GenScript (Table S1), and the PrimerSTAR HS polymerase was utilized for gene amplification through PCR. Based on strain IMX581, the codon-optimized gene ERG1 from *K. marxianus* (kmERG1) was integrated to replace the native ERG1 (scERG1) using CRISPR/cas9, yielding HL01. All the design and construction of the plasmid follows the previously described method^61^. The gRNA cassette for target gene scERG1 was obtained using the single-stranded oligos gRNA-ERG1-F/ gRNA-ERG1-R, followed by assembling with the linearized backbone plasmid pMEL10, the single gRNA plasmid was constructed by Gibson assembly. The repair fragment containing kmERG1 with round 60bp overlap was amplified by primers kmEGR1-scERG1up-F/ kmEGR1-scERG1dn-R using codon-optimized kmERG1 as template. Then the repair fragment and single gRNA plasmid were co-transformed into IMX58. All the strains and primers used in this study were listed in Tables S2 and S3.

#### Strain Cultivation Under Different Temperatures

The thermotolerance was tested and compared between *S. cerevisiae* IMX581 and HL01. Five single colonies of each strain were selected and pre-cultured in YPD media at 30 °C, and cells were then transferred to flasks in 20 mL YPD media to reach 0.1 initial OD600 cultured at 40 +/- 0.1 °C, 200 rpm. After that, the cells were transferred into fresh YPD media every 24h with 0.1 initial OD600 and cultivated at 40 +/- 0.1 °C, 200 rpm.

## Software and Code availability

All simulations of genome-scale models were carried out with Cobrapy^61^ with Gurobi (Gurobi Optimization, LLC) solver. All code is available on Github (https://github.com/Gangl2016/GETCool).

## Author contributions

GL and JN conceptualized the project; GL designed and performed all computations; GL, JZ, BJ, HW, AZ and JN interpreted results. YH performed experimental validations; GL, JZ, HW and YH wrote the initial draft manuscript. All authors carried out revisions on the initial draft and wrote the final version.

## Acknowledgements

The authors would like to thank Tyler W. Doughty, Benjamín J. Sánchez, Avlant Nielsen and Ibrahim Elsemman for the helpful discussions. GL and JN have received funding from the European Union’s Horizon 2020 research and innovation program under the Marie Skłodowska-Curie program, project PAcMEN (grant agreement No 722287). JN also acknowledges funding from the Novo Nordisk Foundation (grant no. NNF10CC1016517), the Knut and Alice Wallenberg Foundation. JZ and AZ are supported by SciLifeLab funding. The computations were performed on resources at Chalmers Centre for Computational Science and Engineering (C3SE) provided by the Swedish National Infrastructure for Computing (SNIC).

