## Supplementary file for "Bayesian genome scale modelling identifies thermal determinants of yeast metabolism"

#### Affiliations

### Supplementary Figures

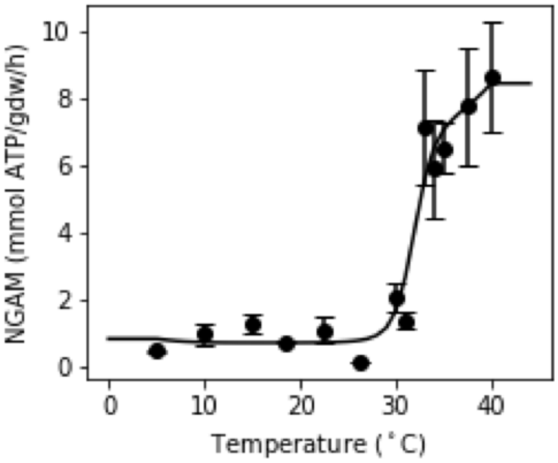


**Fig S1**. Fit an empirical equation to describe the temperature dependence of non-growth associated ATP maintenance (NAGM). The experimental data was collected from Zakhartsev M. *et al.* [1]. The line represents the fitted curve: $NGAM(T) = 0.740 + \frac{5.893}{1+e^{31.920-(T-273.15)}} + 6.12 \times 10^{-6} \times (T - 273.15 - 16.72)^4$.

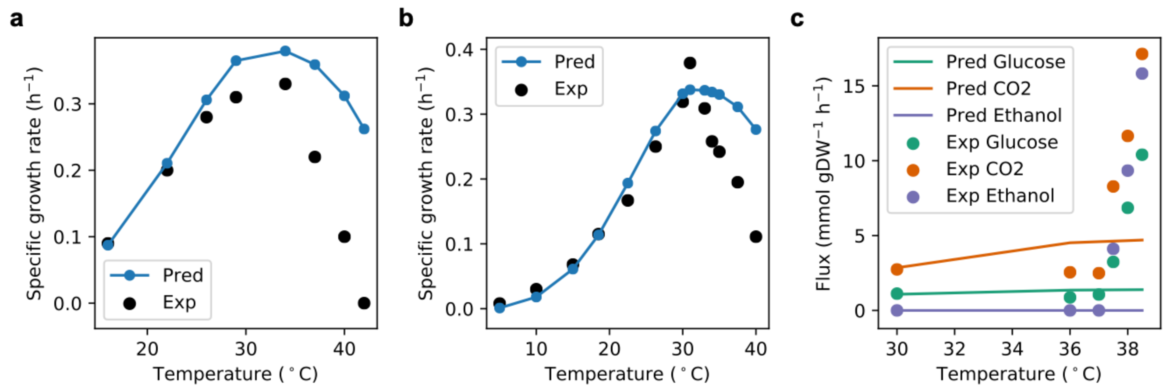


**Fig S2**. Simulated results of specific growth rate at (a) aerobic, (b) anaerobic batch cultivations and (c) fluxes at chemostat cultivation, with the model equipped with initial parameters as described in Methods M5. The black dots are experimental data collected from Caspeta L. *et al*. [2] for (a), Zakhartsev M. *et al*. [1] for (b) and Postmus J. *et al*. [3] for (c).

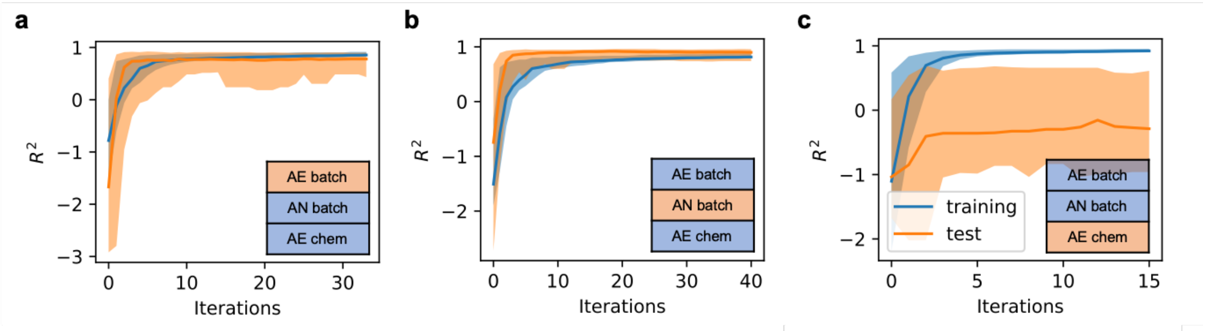


**Fig S3**. Three-fold cross-validation of SMC-ABC approach. Figure legends: AE batch and AN batch are the specific growth rate datasets in aerobic [4] and anaerobic [1] batch cultivations, respectively, and AE chem are the experimental flux measurements in aerobic chemostat cultivation [3]. For each fold, the two datasets (blue) were used to update the *Prior* model and were then tested on the remaining dataset (orange) during iterations. All the *Posterior* models showed an improved performance on the test dataset through iterations. Whereas *Posterior* models trained on both batch and chemostat data showed good performance on the rest of the batch data (aerobic or anaerobic, Figure 2b,c), ones trained on batch data (aerobic and anaerobic growth rate) showed poor performance on chemostat data (Figure 2c: $R^2_{test} < 0$). These findings indicated it is necessary to use all datasets to update the *Prior,* especially since batch and chemostat data provide non-overlapping, orthogonal information.

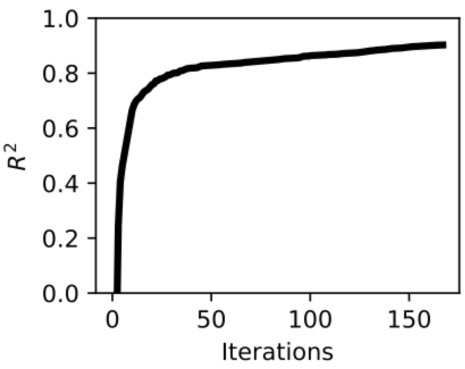


**Fig S4**. The average $R^2$ score on three datasets during iterations in the Sequential Monte Carlo based Approximate Bayesian Computation (SMC-ABC) approach.

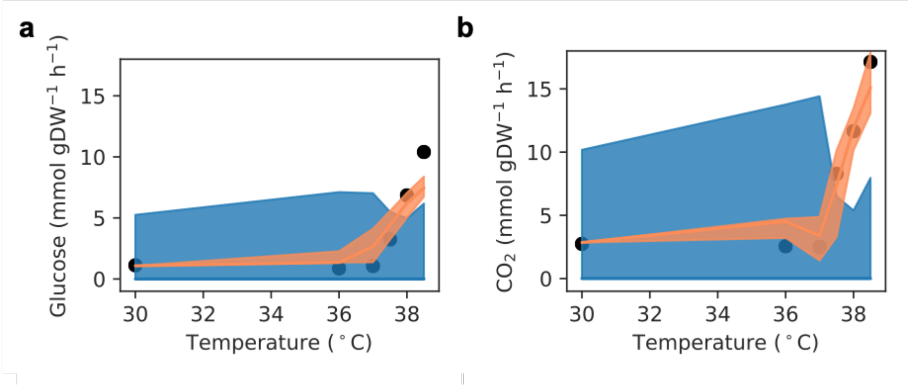


**Fig S5**. Simulated (a) glucose uptake and (b) $CO_2$ secretion fluxes at various temperatures with *Posterior* models. Lines indicate median values and shaded areas indicate regions between the 5-th and 95-th percentiles. Black dots show the experimental values from Postmus J. *et al*. [3]

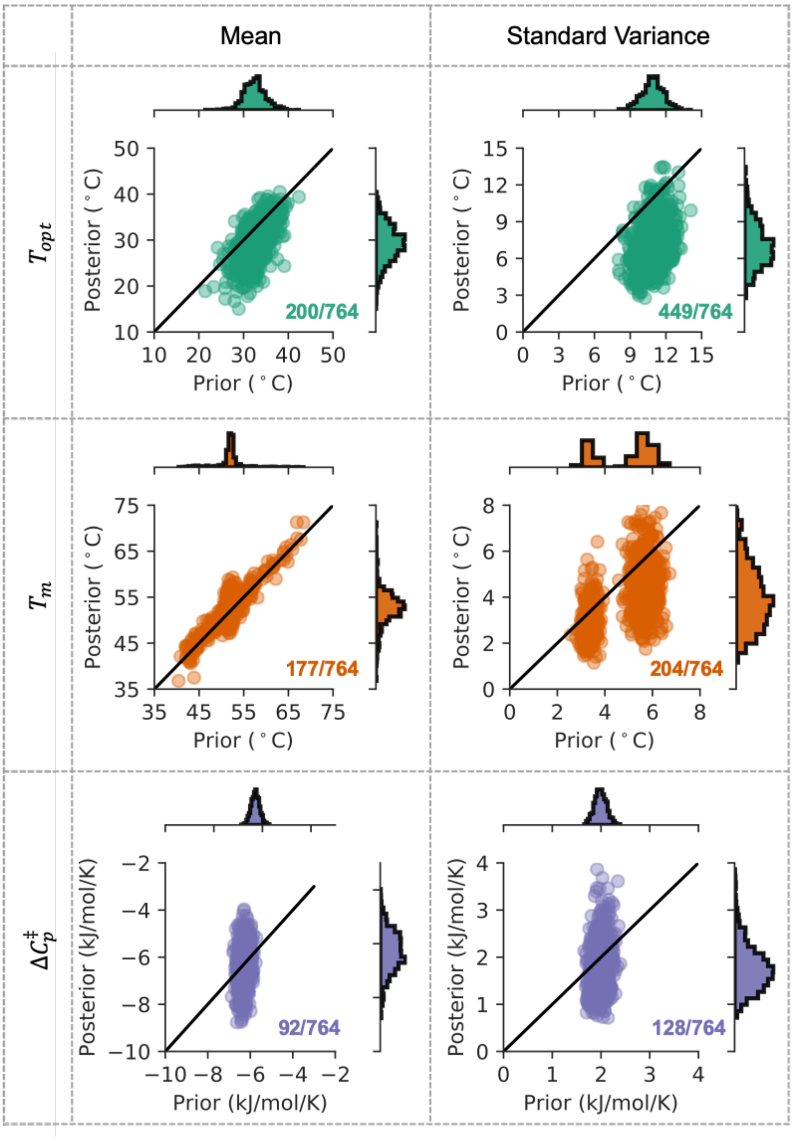


**Fig S6**. Parity plots of updated mean and standard variance of enzyme $T_{opt}$s , $T_m$s and $\Delta C_p^{\ddagger}$s. The mean and standard variance are calculated based on 128 *Prior* and 100 *Posterior* models. The inset numbers indicate the number of enzymes, out of all 764, with a significantly changed mean (Šidák adj. Welch's *t*-test *p*-value < 0.01) and variance (Šidák adj. one-tailed *F*-test *p*-value < 0.01).

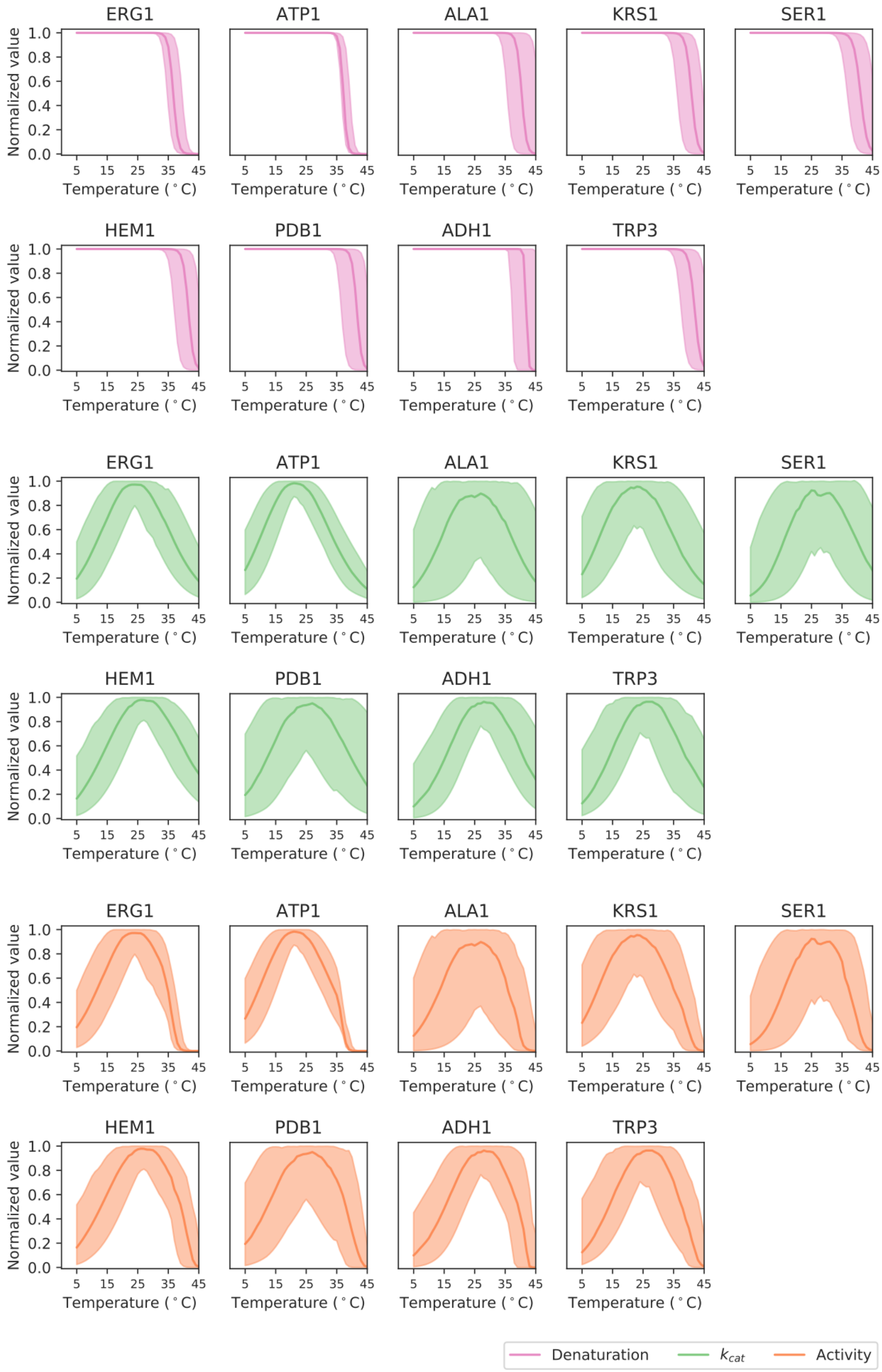


**Fig S7**. The temperature effect on 9 enzymes with a mean melting temperature below 42 °C in the 100 *Posterior* models. The denaturation is shown as the probability of an enzyme in the native state. The kcat is shown as normalized value by the maximal kcat. Activity, the specific activity of an enzyme is shown as the product of values in denaturation plots and kcat plots. Lines indicate median values and shaded areas indicate regions between the 5-th and 95-th percentiles.

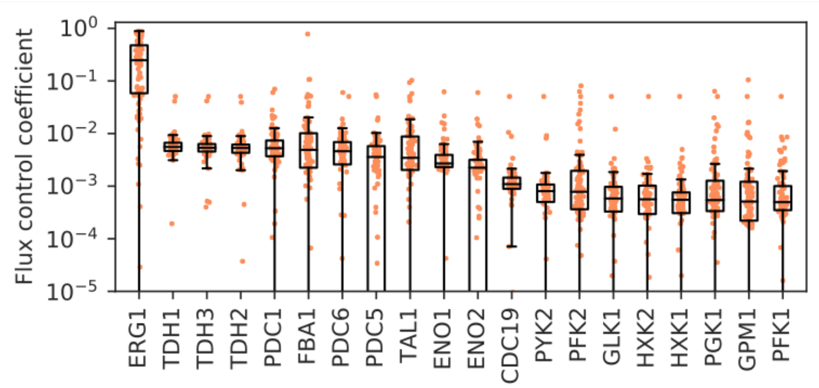


**Fig S8**. 20 enzymes with the highest flux sensitivity coefficients at 42 °C.

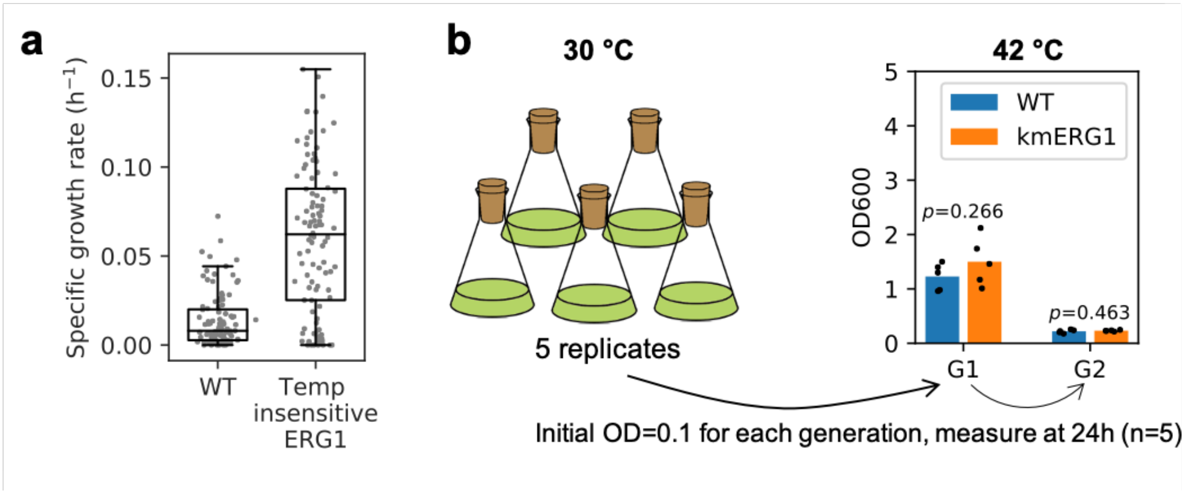


**Fig S9**. ERG1 at 42 °C. (a) Predicted maximal specific growth rate of wide-type yeast and the one without any temperature constraints (fully functional) on ERG1 enzyme at 42 °C. (b) The effect of KmERG1 expression on thermo tolerance in *S. cerevisiae*. The strains were cultivated at 40 °C for six generations to reach the steady state of growth. Optical densities (600 nm) are shown at 24 h. Each bar indicates the mean and dots represent the values of 5 replicates. *p*-values denote Welch's *t*-test.

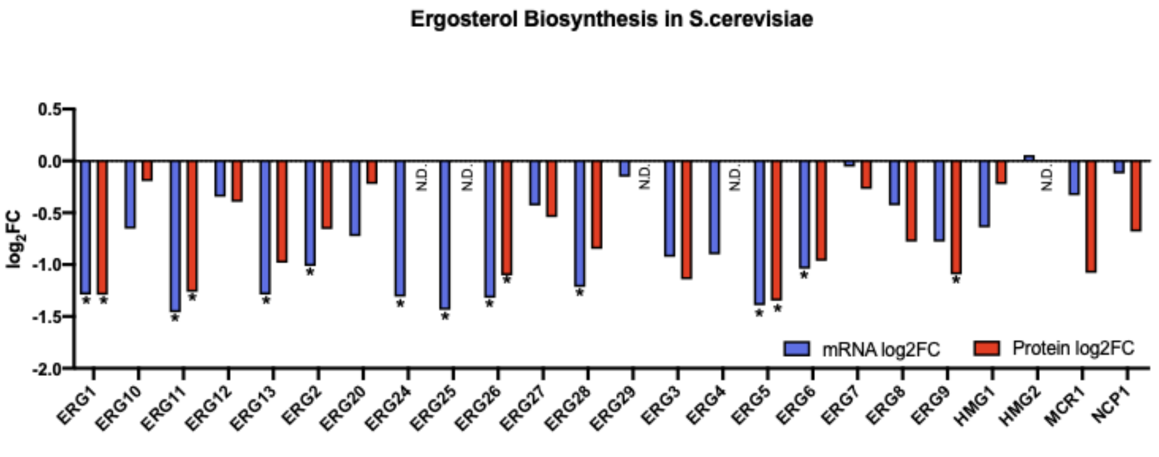


**Fig S10**. Downregulated ergosterol biosynthesis pathway at both transcription and translation levels. The proteomics and transcriptomics data were from Doughty T. *et al.*[5]. * indicates the genes with an absolute log2FC>1 and FDR<0.01.

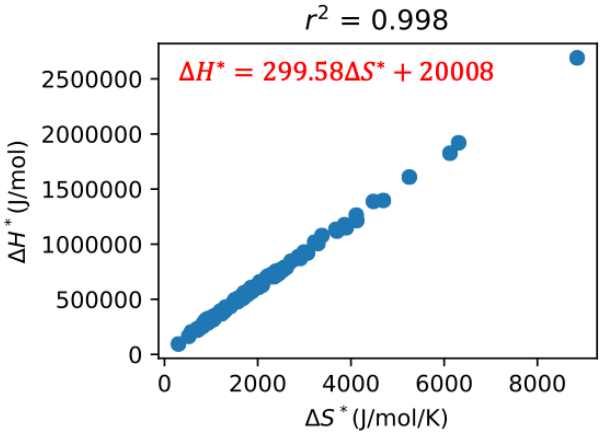


**Fig S9**. The correlation between enthalpy and entropy changes of protein denaturation process at convergence temperatures (373.5 K for $\Delta H^*$ and 385 K for $\Delta S^*$). The plot shows data of 116 proteins from Sawle L et al [6].

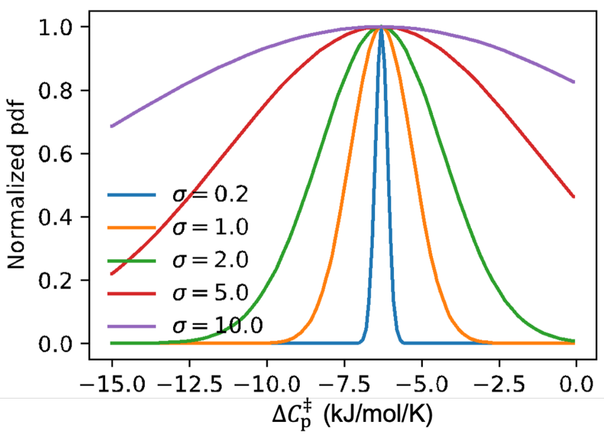


**Fig S10**. The normal distributions of $\Delta C_p^{\ddagger}$ with a mean of -6.3 kJ/mol/K and different standard variances.

### Supplementary Tables

**Table S1**. Codon-optimized kmERG1

| | **Sequence (5’-3’)** |
|---|---|
| *kmERG1* | ATGTCTTCAGCTACTGATAAGAAAGTTGTTCCACAAGAATTGTTGAATGCTGATGCATCTGTTGTTTATG<br>ATGCAATTATTGTTGGTTGTGGTGTTGTTGGTCCATGTATTGCTACTGGTTTAGCAAGAAAGGGTAAAAA<br>GGTTTTGATCGTTGAAAGAGATTGGACAATGCCAGATAGAATTGTTGGTGAATTAATGCAACCAGGTGG<br>TTTGAGAGCTTTAAGATCATTGGGTATGATCCAAGCTATTAATAACATCGATGCAATCCCAGTTACTGGT<br>TACTCTGTTTTCTATAACGGTAAAGAAGTTTCAATCCCATACCCTTTTAAAGCTGATGTTGCACCAGTTG<br>AAAAGATTCCAGATTTGGTTTTCGATGGTAACGATAAGGTTTTGGAAGATAACACTATCCATGTTAAGGA<br>ATTTGAAGATGGTGAAAGAGAAAGAGGTGTTGCTTTTACACATGGTAGATTTTTACAAAATTTGAGAGAT<br>ATGGTTTCTAAAGAAAAGAATGTTACAAGATTACAGGGTAACGTTATCGAAATTTTGAAAAATAAGGAAA<br>ATGAAGTTGTTGGTGCTAAAGTTGATATTCCAGGTCGTGGTAAAGAAGAGTTTAAAGCATATTTGACTTT<br>CGTTTGTGATGGTATTTTCTCTCATTTCAGAAAGGAATTGGCTTCAGATCATATTCCAACAGTTGGTTCT<br>TCATTTGTTGGCATGTCTTTGTTTAATGCTGATGTTCCAGCTAAAAATCATGGTCATGTTATCTTGGGTA<br>CTAATCACATGCCAGTTTTGGTTTACCAAATTTCTCCAGAAGAAACAAGAATTTTATGTGCTTACAACTC<br>ACCAAAATTGCCAAAGGATATTAAGTCTTGGTTGAAGTCAGATGTTAGACCAAATTTGCCAAAATCTTTG<br>TTGCCATCATTTGATAAGGCAGTTGAAGATGGTAAATACAGATCAATGCCAAATTCATACTTACCAGCTA<br>AGCAAAACACTATCACAGGTTTGTGTGTTATTGGTGACGCATTAAATATGAGACATCCATTGACTGGTG<br>GTGGTATGGCTGTTGGTTTGAACGATGTTGCATTGTTGATTAAAACTATCGGTAATTTGGATTTCTCTGA<br>TAGAGAAACAGTTTTGGATGAATTGTTAGAATACCATTACGAAAGAAAGTCTTATGATTCAGTTATTAAT<br>GTTTTGTCTATTGCTTTATTTTCATTGTTCGCTGCAGAAAATAAGAATTTGCAAGTTTTGCAAAGAGGTTG<br>TTTCAAGTACTTCGAAAGAGGTGGTGACTGTGTTAACATCCCAGTTTCATTTTTAGCAGGTGTTATGCCA<br>AAGCCATTTTTGTTGACTAAGGTTTTCTTTGCTGTTGCATTGTACTCTATCTATGTTAACTTCCAAGAAAG<br>AGGTATCGTTGGTTTTCCATTAGCTGTTATTGAAGCAATTTCAATCTTGATTACTGCTGCAAGAGTTTTTA<br>CACCATATTTGTTTAGAGAATTGACAGGTTAA |

**Table S2**. List of strains.

| Strain name | Genotype | Reference |
|---|---|---|
| IMX581 | *MATa ura3-52 can1Δ::cas9-natNT2 TRP1 LEU2 HIS3* | [7] |
| HL01 | *MATa ura3-52 can1Δ::cas9-natNT2 TRP1 LEU2 HIS3 ERG1Δ::kmERG1* | This study |

**Table S3**. List of primers

| Primer name | Sequences (5´-3´) |
|---|---|
| tCYC1-X-2dn-R | GATAAATCTTCAGCATAGATGGGTAACGGGATCCCTCTGTGAGGGCCGATTATGCAGGCCT AGACCCGGCCGCAAATTAAAGCCTTCGAG |
| gRNA-ERG1-F | GTTGATAACGGACTAGCCTTATTTTAACTTGCTATTTCTAGCTCTAAAACAATGTTACTAGAG TGCAAGGGATCATTTATCTTTCACTGCGGAGAAGTTTCGAACGCCGAAACATGCGCA |
| gRNA-ERG1-R | TGCGCATGTTTCGGCGTTCGAAACTTCTCCGCAGTGAAAGATAAATGATCCCTTGCACTCTA GTAACATTGTTTTAGAGCTAGAAATAGCAAGTTAAAATAAGGCTAGTCCGTTATCAAC |
| kmEGR1-scERG1up-F | CAATTGTCCAGTATTGAACAATACAGGTTATTTCGAACAATTGAAAAAAAAAAATCACAGAAA AACATATCGAGAAAAGGGTCATGTCTTCAGCTACTGATAAGAAAG |
| kmEGR1-scERG1dn-R | GCCTTCCAAGCCGACTTCTGTAATAAAAAAAAAAAGGTGCAGCTTAATGTTTGACGGTTCCTA TCCTCTCTCCCTTATAAGCTGTAGTTAACCTGTCAATTCTCTAAAC |
